# Young adults accelerate their arms significantly faster and earlier than old adults resulting in improved center of mass dynamics during an overground slip perturbation

**DOI:** 10.1101/2023.12.09.570848

**Authors:** Jonathan S. Lee-Confer, Matthew K. Lo, Karen L. Troy

**Affiliations:** University of Arizona, Department of Physical Therapy, Tucson, AZ, USA; Verum Biomechanics, Tucson, AZ, USA; University of California, Irvine, Irvine, CA, USA; Worcester Polytechnic Institute, Worcester, MA, USA

**Keywords:** aging, falls, balance, gait, stability, posture

## Abstract

Arm motions reduce the likelihood of falling from a slip in young adults, and injuries to older adults from falls are known to occur from a frontal plane fall. Therefore, the purpose of this study was to quantify the frontal plane trunk and arm motion in older and younger adults during a slip. Eleven older adults (age: 72.0 ± 5.0) and 11 younger adults (age: 25.17 ± 5.94) were subjected to a slip during walking. 27% of the older adults and 11% of the younger adults experienced a fall from the slip incident. The arm excursion did not significantly differ between groups, however younger adults’ average peak arm abduction occurred 310 ms prior to older adults (p = 0.03). There were no significant differences in peak trunk between groups. The arm abduction acceleration was significantly higher in younger adults compared to older adults (3593.2 ± 1144.8 vs. 2309.8 ± 1428.5 degrees/s^2^, p = 0.03). Furthermore, young adults exhibited a significantly reduced CoM excursion compared to old adults (4.55 ± 3.45 vs. 10.27 ± 6.62 cm, p < 0.01). Young adults may have exhibited lower fall rates due to the significantly higher acceleration of their arms compared to old adults.

## Introduction

One of the largest cause of injuries to adults over the age of 65 are from falls and approximately 25% of adults over the age of 65 will fall on any given year (1,2). The projected annual cost on the United States’ health care system for older adults falling is expected to be $100 billion by the year 2030 (3). Furthermore, it is reported that 56% of falls in older adults occur from stepping on a slippery surface and experiencing a slip (2). The majority of the biomechanical research on slip perturbations have focused on the lower extremities (4,5,14–23,6–13). There are fewer investigations of the upper extremities during a slip incident as a consequence of focusing on the lower extremity responses (24–30). It is important to investigate how older adults move their arms as it is shown that moving the arms reduces the likelihood of falling by 70% (31). As such, understanding how older adults move their arms during a slip and characterizing strategies that increase the likelihood of recovery is a key prerequisite to developing interventions to reduce falls in this vulnerable population.

Slip perturbations induce a sideways loss of balance and require extremity movements to restore stability. It is reported that younger adults exhibited up to 54 degrees of lateral trunk flexion during a slip incident (32). Excessive lateral trunk flexion can be problematic for older adults as hip fractures are known to occur from a sideways loss of balance (33–35). During a slip, the perturbed foot slides anteriorly and the trailing leg is necessary for regaining balance (36). The lower extremities are likely unable to assist in controlling frontal plane stability when a sideways loss of balance begins to occur during a slip as the legs are focused on restoration of the base of support. The upper extremities have been reported to produce three times higher excursion in the frontal plane than the sagittal plane during a slip incident (24) and therefore, the arms are more likely to be able to control a sideways loss of balance. It is important to understand the frontal plane arm mechanics that can improve balance within the frontal plane to reduce the likelihood of a sideways fall in older adults.

Within the paradigm of the upper extremities, most studies have focused on sagittal plane motions during a slip incident. Arm motions were first described to undergo arm flexion in response to a slip perturbation (25). The motion of the arms in the sagittal plane were further described in other studies (26–28). Furthermore, these studies reported that the arm motions during a slip assisted in controlling the center of mass, reducing trunk extension velocity, maintaining whole-body angular momentum, or even used for grasping nearby objects for balance. However, the utility of the arms within the sagittal plane in preventing serious injuries in older adults may be limited, as sagittal plane motion has little contribution to sideways balance.

The research on the frontal plane motion of the arms during a slip perturbation is limited. One study has reported that frontal plane excursion of the arm contralateral to the slipping foot during a slip incident is nearly three times larger compared to the sagittal plane excursion (24). Another study reported that falls were reduced by 70% when individuals swung their arm contralateral to the slipping foot during a slip perturbation (31). The mechanism of frontal plane arm motion was reported to reduce lateral center of mass excursion by 37.5% during a slip incident (37). Currently there are no studies investigating older adults’ frontal plane motion of the arms during a slip and this is an important area to study as this may lead to fall reduction.

Therefore, the purpose of this study is to characterize and compare the frontal plane motion of the contralateral arm to the slipping foot and frontal plane trunk excursions, and the frontal plane center of mass dynamics in older and younger adults. We hypothesize that younger adults will exhibit significantly increased frontal plane arm excursion, significantly greater acceleration of arm movement responses, and decreased lateral trunk excursion and decreased center of mass excursion compared to older adults.

## Methods

### Participants

A subset of 22 participants from a larger study investigating the biomechanical mechanisms of slip recovery were analyzed for this study (38). Participants were analyzed in this study if they exhibited a minimum of 10 degrees of lateral trunk flexion. As such, eleven community-dwelling older adults and eleven younger adults were analyzed in this IRB approved study. Participants’ anthropometrics for this study may be found in Table 1. All participants signed a written informed consent after being provided with the scope of this study. Although participants knew they would be slipped, they did not know when or how this would take place. All participants were screened by an attending physician and were excluded if they exhibited any musculoskeletal, cardiovascular or neurological conditions.

**Table 1.** Participants’ demographic and anthropometric information.

| Participants | Age (years) | Height (in.) | Weight (lbs.) | Sex (M/F) |
| --- | --- | --- | --- | --- |
| Older adults (n=11) | 72.0 ± 5.0 | 66.36 ± 4.52 | 160.55 ± 32.69 | 6/5 |
| Younger adults (n=11) | 25.45 ± 6.14 | 69.82 ± 4.67 | 168.55 ± 41.68 | 8/3 |

### Instrumentation

Detailed experimental methods have been previously described (38). Briefly, all walking trials were performed on a designated walkway within the laboratory at the University of Illinois at Chicago. A 1.22 m x 2.44 m x 0.63 cm Plexiglas sheet was imbedded into the walkway. Participants walked several times across the walkway before being slipped. During the slip trial, a thin film of dried water-soluble lubricant was quickly and quietly activated with a spray bottle of water at a time unbeknownst to the older participants. Younger adults were subjected to an unexpected slip perturbation induced by stepping onto a thin applied layer of mineral oil.

Motion analysis was modeled in three dimensions with an eight-camera motion capture system collecting data at 60 Hz (Motion Analysis, Santa Rosa, CA). Reflective joint markers were carefully secured to anatomical locations that were utilized to produce a 13-segment rigid body model to generate joint kinematics (OrthoTrak, Motion Analysis, Santa Rosa, CA).

Fall-prevention equipment was utilized to prevent injuries occurring to participants in case an actual fall were to occur. All participants were fitted with a safety harness that supported the entire body weight of the participant and prevented contact of the participants’ hands, knees, or buttocks with the floor if a fall were to occur. Individuals were allowed to wear their own shoes that they were comfortable with.

### Procedures

Participants were allowed to practice walking on the walkway without being attached to the safety harness to gain experience walking in an unfamiliar environment. The participants were aware that may be slipped on any trial, however the participants were not aware of how many control trials would take place, nor the which trial the transition of the surface from dry to wet would occur at. Furthermore, the participants were unaware of the mechanism that would induce a slip. Each participant was exposed to only one slip and the slip was randomly selected to occur on either the left or right foot.

### Data Analysis

Slip onset was defined as the instantaneous moment of weight acceptance onto the slippery surface. A slip trial was considered to be a fall if the participants’ CoM dropped below 95% of their minimum vertical CoM during gait (39). The peak frontal plane angle of the arm contralateral to the slipping foot was calculated as previous work suggested the arm contralateral to the slipped foot produced the frontal plane motion of interest (24). Peak acceleration of the arm contralateral to the slipping foot, and peak frontal plane trunk flexion angles were calculated. Frontal plane CoM excursion was calculated by taking maximal lateral distance away from the CoM frontal plane position at heel strike. To characterize the behavioral differences between younger and older adults, custom-written MATLAB (Mathworks, Natick, MA, USA) code was created to conduct a statistical parametric mapping test to determine if there were significant differences in compensatory acceleration responses at time points along the continuum after a slip perturbation was initiated. One-tail independent t-tests were conducted in SPSS 16.0 software (SPSS, Chicago, IL, USA) to determine if younger adults’ peak arm abduction was higher, peak arm acceleration was lower and center of mass excursion was lower within the frontal plane compared to older adults. A Pearson correlation coefficient test was conducted to determine if there was a relationship between peak trunk flexion angles and peak contralateral frontal plane arm excursion. All statistical tests used an alpha criterion value of 0.05 to determine significance.

## Results

Twenty-seven percent of the older adults (n = 3) and 11% of the younger adults (n = 1) experienced a fall from the slip incident. Using a t-test outlier test, one older adults’ data for time to peak abduction of 2.1 seconds resulted in a Z-score of +2.44 (critical boundary was +2.35), was recorded as an outlier and the data from this participant was not included in the analyses. The removal of this data did not affect significance of any variables measured in this study. As such, the overall data analysis included 11 young adults and 10 older adults. The frontal plane excursion of the contralateral arm to the slipping foot did not significantly differ between younger and older adults (71.97 ± 28.16 vs. 83.32 ± 25.05 degrees, p = 0.34, Fig. 1), however older adults were significantly delayed in reaching peak arm abduction compared to younger adults (853 ± 509 vs. 542 ± 67 milliseconds, p = 0.03). There were no significant differences in frontal plane lateral trunk flexion between older and younger adults (16.1 ± 4.5 vs. 17.03 ± 5.28 degrees, p = 0.66). The average peak arm abduction acceleration in the contralateral arm was significantly higher in younger adults compared to older adults (3593.21 ± 1144.80 vs. 2309.83 ± 1428.48 degrees/s^2^, p = 0.03, Fig. 2). Furthermore, in the contralateral arm to the slipping foot, young adults exhibited significantly higher acceleration of arm abduction than older adults between 133 ms and 200 ms following the foot contacting the slippery surface (p < 0.05, Fig. 2). Young adults significantly reduced their frontal plane CoM excursion compared to old adults (4.6 ± 3.5 vs. 10.47 ± 6.6 cm, p < 0.01, Fig. 3). Lastly, a significant moderate correlation was found between peak lateral trunk flexion and peak frontal plane arm excursion (r = 0.52, p < 0.02, Fig. 4). An example of an individual slipping and exhibiting contralateral arm abduction and trunk lateral flexion is shown in figure 5.

**Figure 1.**
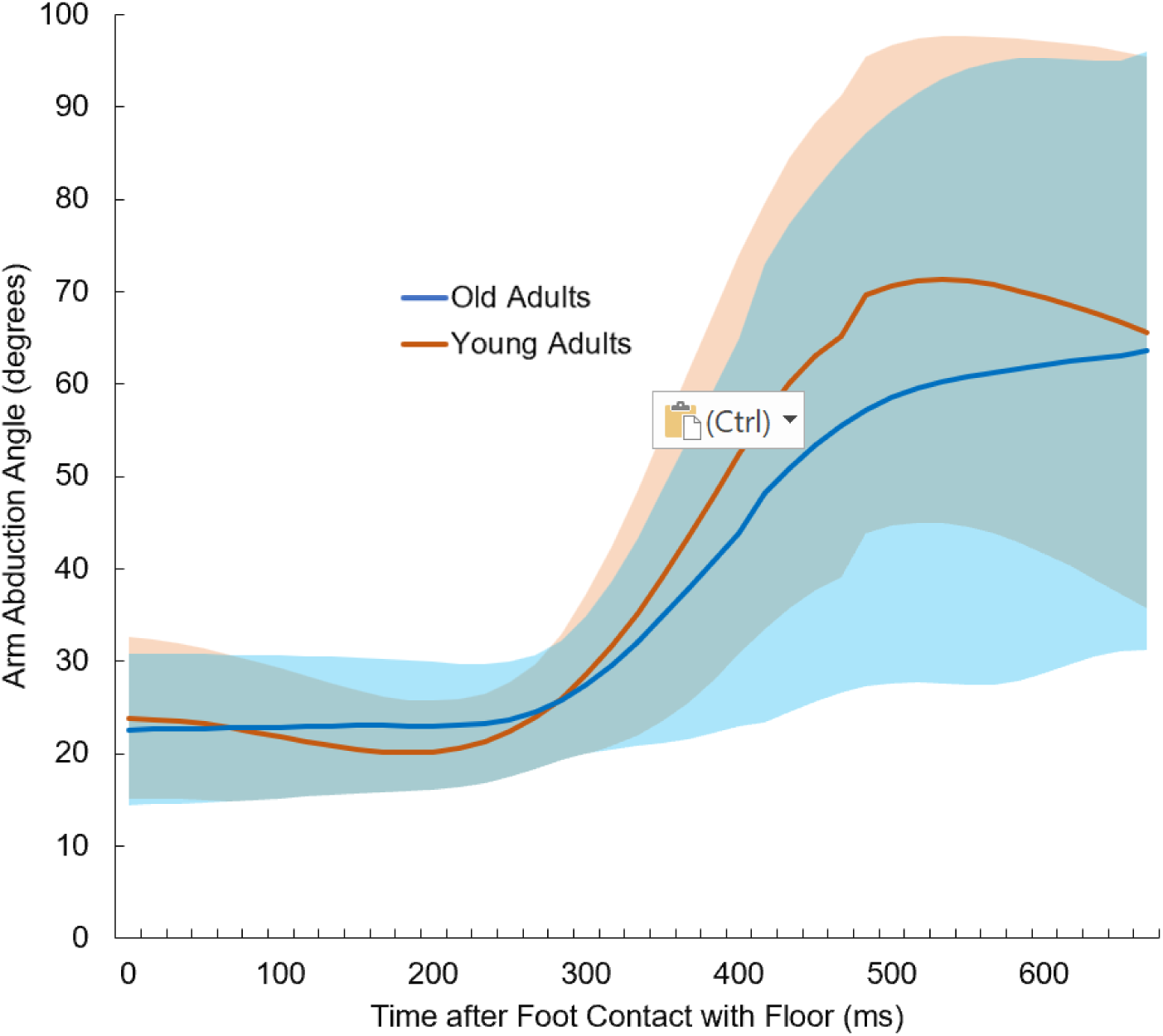
Contralateral arm to the slipping foot arm abduction angles of younger adults (orange line, n = 11) and older adults (blue line, n = 10). Peak average arm abduction of younger adults occurred 310 ms prior to older adults (p < 0.02).

**Figure 2.**
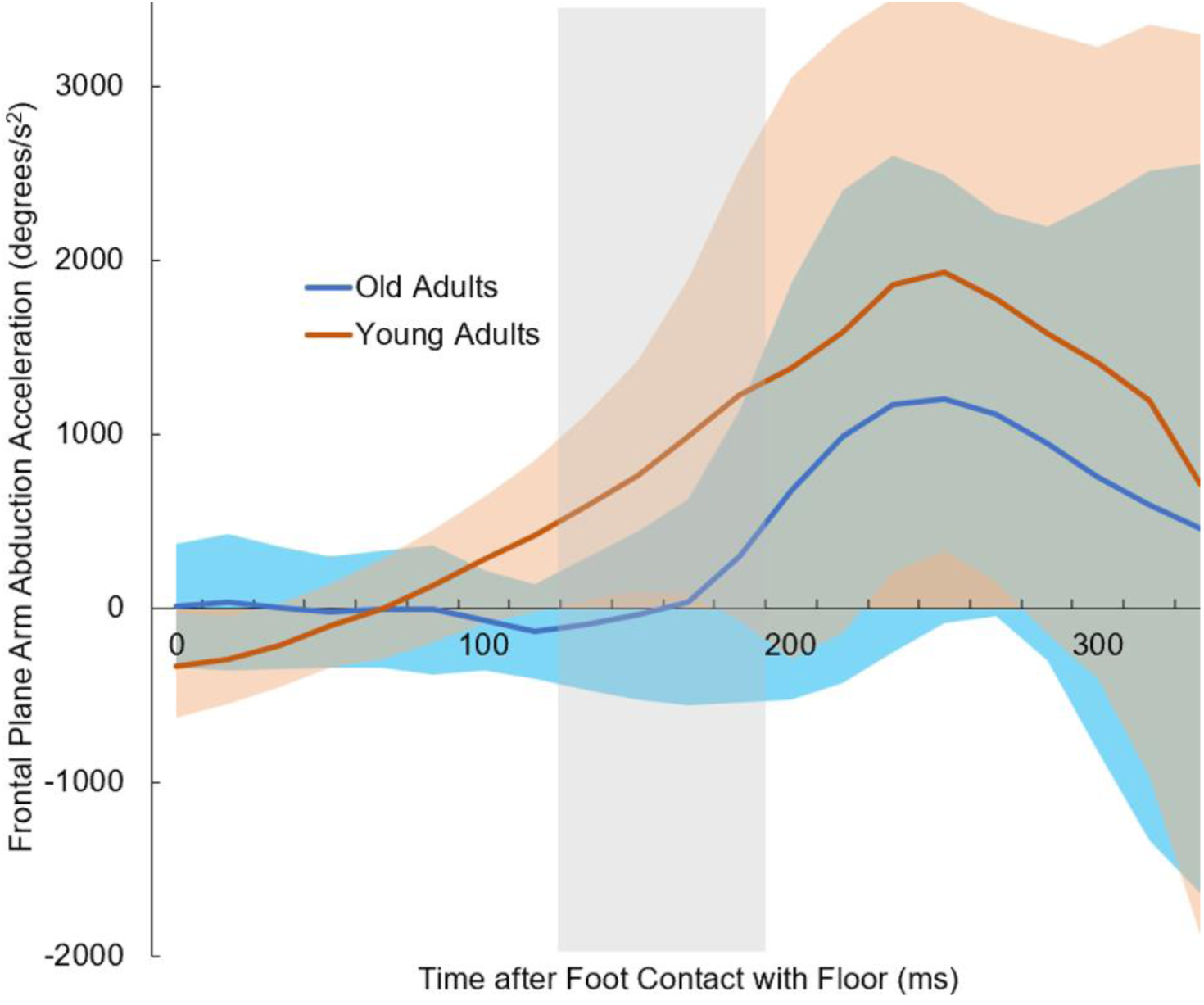
Frontal plane arm acceleration of younger (n = 11) and older adults (n = 10) during a slip perturbation. The blue line represents the average arm acceleration of the older adults and the orange line represents the average arm acceleration of the younger adults. The shaded gray region highlights 133-200 ms after the foot contacted the slippery floor and designates significant differences between the younger and older adults as reported by statistical parametric mapping (p < 0.05).

**Figure 3.**
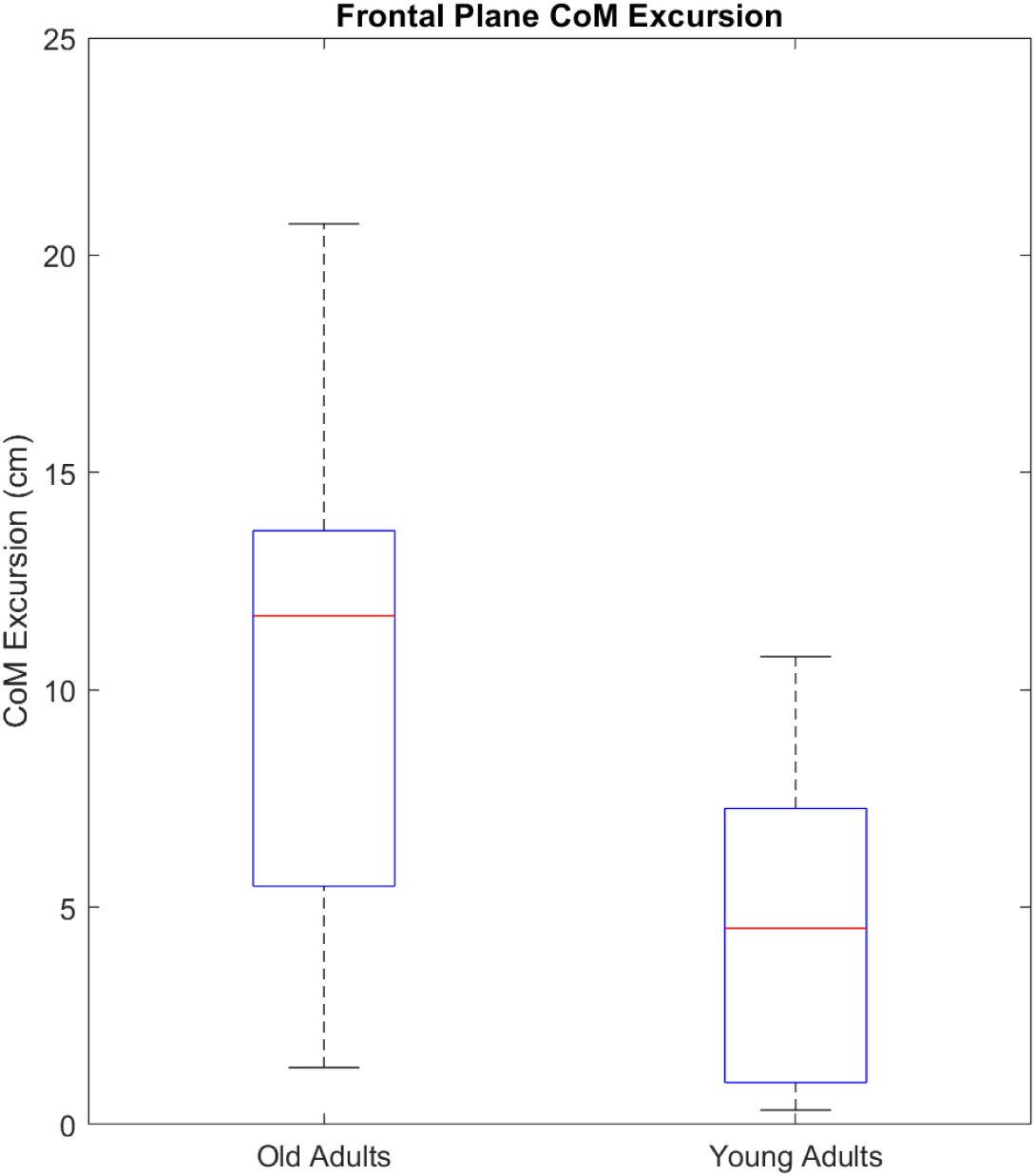
Boxplots showing frontal plane CoM excursion between old (n = 10) and young adults (n = 11). Young adults significantly reduced their frontal plane CoM excursion compared to old adults during a slip incident (p < 0.01). The horizontal red line represents the median for each group.

**Figure 4.**
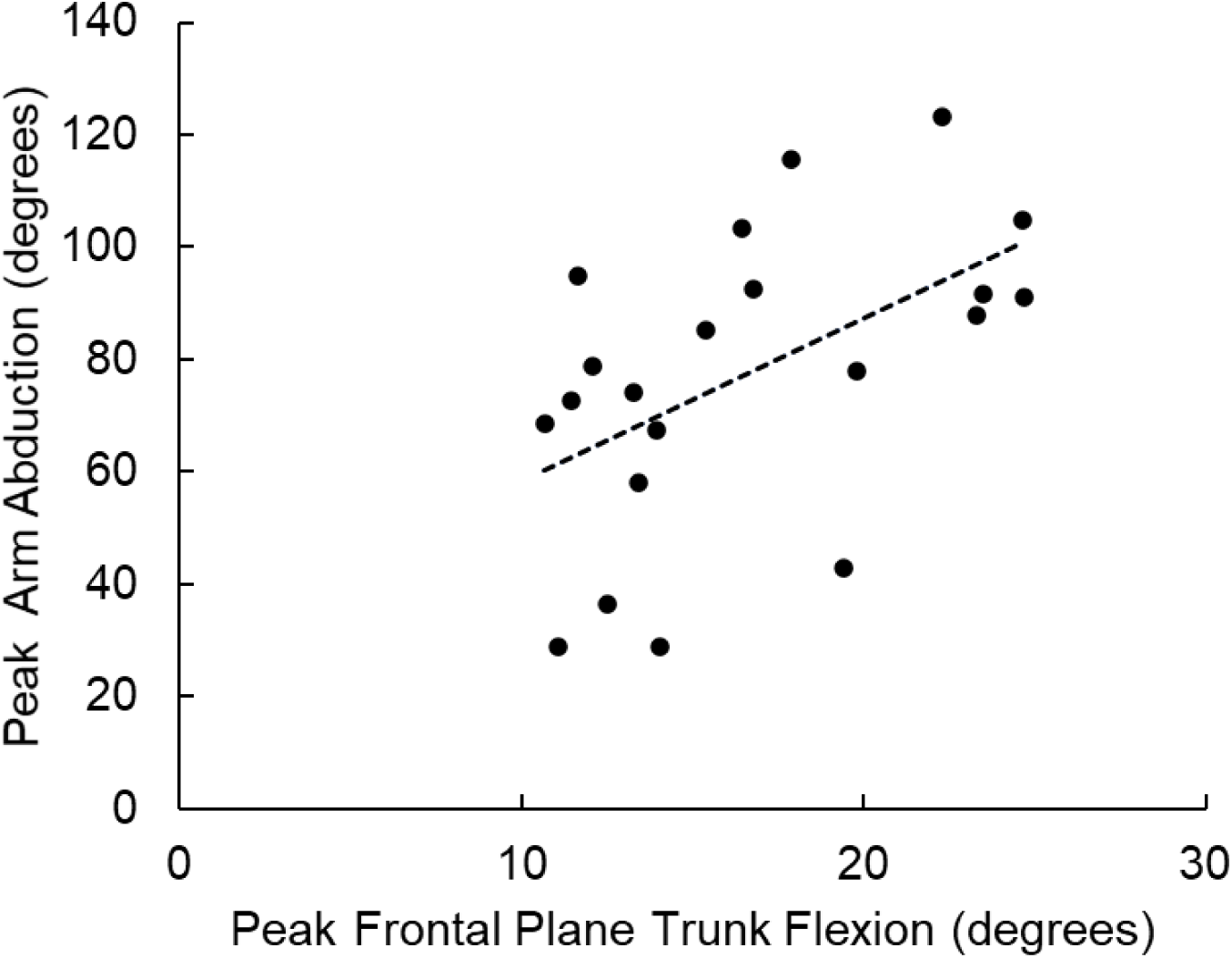
A graph depicting the correlation between peak frontal plane trunk flexion and peak arm abduction of young and old adults (r = 0.52, p < 0.02, n = 21).

**Figure 5.**
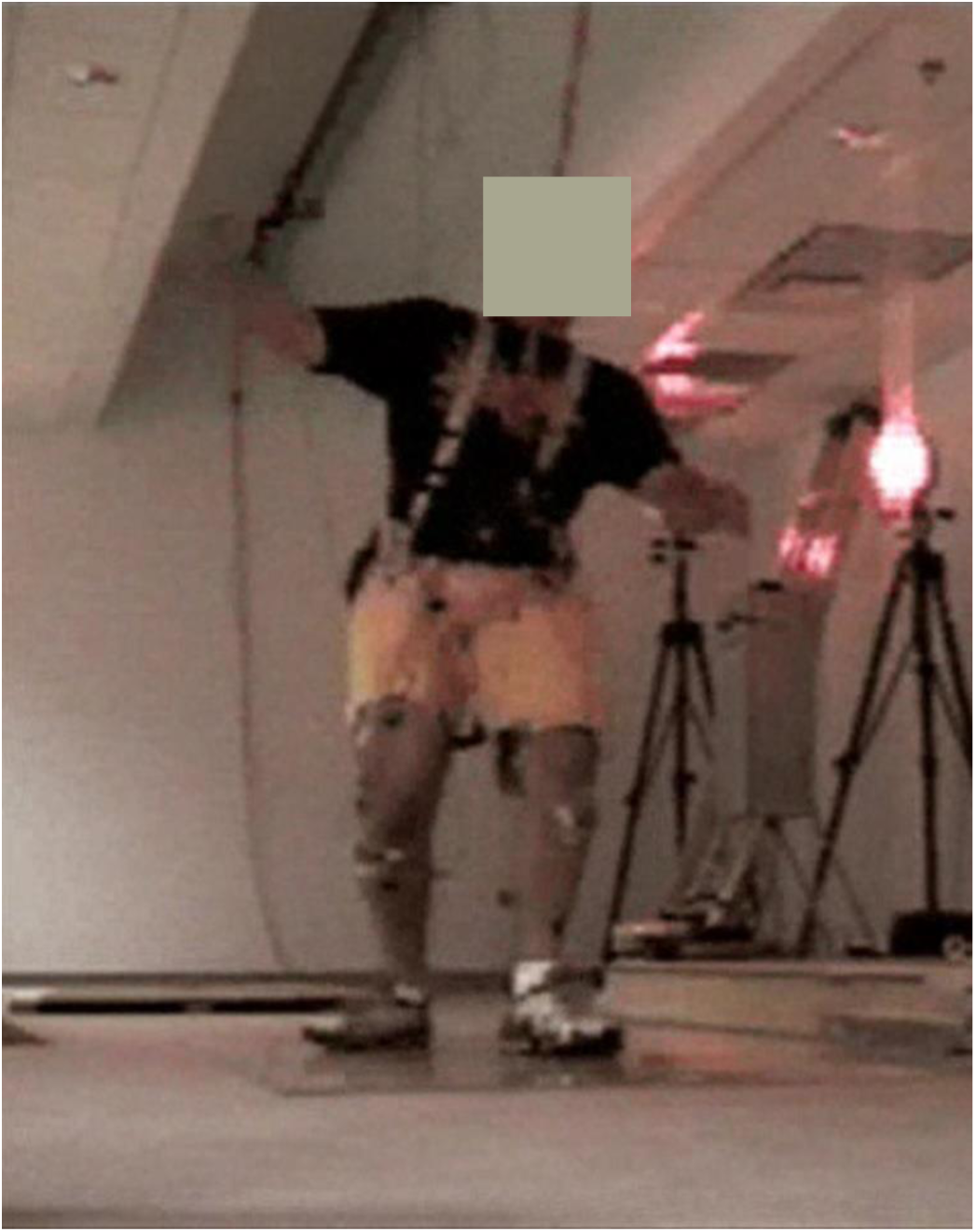
A photograph of a participant experiencing a slip on their left foot and exhibiting left trunk flexion and right arm abduction.

## Discussion

The purpose of this current study was to characterize and compare the frontal plane motion of the arm contralateral to the slipping foot and the frontal plane arm acceleration of older adults and younger adults during a slip perturbation. Contrary to our hypothesis, older adults exhibited a similar amount of frontal plane arm excursion compared to younger adults during a slip incident. In support of our hypothesis, older adults exhibited significantly slower arm acceleration within the frontal plane and increased lateral center of mass excursion compared to younger adults during a slip perturbation.

The findings of total frontal plane arm excursion in this study were similar to a previous study reporting the frontal plane motion of the contralateral arm to the slipping foot. In a prior study, it was reported that healthy and young adults exhibited 61.7 ± 26.9 degrees of abduction in the contralateral arm to the slipping foot during a slip incident (24). Similarly, this study found that healthy and young adults exhibited 72.0 ± 28.2 degrees of abduction in the contralateral arm to the slipping foot during a slip incident and older adults exhibited 83.3 ± 25.1. The similarity of arm responses between older and younger adults in this study was surprising as a previous study reported significant differences between younger and older adults’ arm movements in the sagittal plane during a slip and reported that younger adults exhibited arm extension moments whereas older adults exhibited arm flexion moments (27). Even though peak arm abduction angles were similar between younger and older adults, older adults’ peak arm abduction occurred 310 milliseconds after younger adults exhibited their peak arm abduction (Fig. 1). It is possible that our study found similar results between younger and older adults because the inclusion criteria required frontal plane trunk motion. While older adults exhibited similar amounts of frontal plane arm excursion compared to younger adults, they differed in the behavioral responses.

Older adults may be more at risk of falling compared to younger adults due to older adults’ inability to generate sufficient arm acceleration. Arm motion during a slip has been shown to regulate angular momentum and reduce slip severity (28), trunk velocity (26), and center of mass excursion by ∼35% and increase margins of stability during a slip perturbation (37). The present findings of older adults exhibiting less acceleration of the arms indicate that older adults may not be utilizing the biomechanical effects of the arms to reduce fall risk during a slip incident. To further illustrate this point, the lateral center of mass excursion was significantly higher in older adults than younger adults. The frontal plane arm acceleration may be the driver of reducing center of mass excursion as older and younger adults exhibited the similar amounts of arm abduction excursion. As the lower extremities serve as the base of support, the legs are unlikely to produce frontal plane movement to counter frontal plane induced trunk mechanics. However, the arms have the capability to produce fall protection mechanisms in all three planes making the study of arm contribution to fall prevention functional and practical. In this study, the delay in peak arm abduction and lower arm abduction acceleration may reduce the protective benefits of the arms to counter frontal plane trunk motion.

From a theoretical biomechanical perspective, an overground slip incident during walking would inherently induce frontal plane motion. A slip incident begins with an individual making foot contact onto a slippery surface where the foot intends on accepting weight. During a slip however, the perturbed foot slides anteriorly away from the body while still maintaining weight on the foot as the slip progresses. The weight bearing on the anteriorly shifting perturbed foot would lower the height of the hip on the perturbed side causing the trunk to rotate within the frontal plane.

Preventing sideways falls is critical to reducing the number of serious injuries in older adults, and slip incidents have been shown to induce sideways motion (32,40–43). A slip during walking would theoretically initiate frontal plane motion. This process starts when a foot contacts a slippery surface under the intention of weight bearing. During the slip, the affected foot slides forward, while supporting weight, causing a drop in the hip height on the same side. This results in trunk rotation within the frontal plane driven by pelvic lateral tilt which has the propensity to expose the hip to impact if a slip results in a fall.

From an observational perspective, the lower extremities predominantly move within the sagittal plane during a slip incident (44). Furthermore, the legs are unable to produce any counterbalance movements against any frontal plane loss of balance as the legs are preoccupied with restoring the base of support. As a consequence, the body’s restoration of balance within the frontal plane relies upon the ability of the upper extremities to produce multidirectional movement.

Previous work has shown that the movements of the arms are effective at reducing center of mass excursion and velocity in the frontal plane, as well as increasing the margins of stability, compared to individuals with their arms constrained (37). Similarly, it is also likely that the arm motion within the frontal plane reduces frontal plane angular momentum and sideways trunk flexion as these reductions are observed in the sagittal plane. Future studies may consider investigating the effect of the arms on whole-body angular momentum of older adults exposed to a slip incident as whole-body angular momentum factors in the velocity of moving segments about the dynamic whole-body center of mass. It is possible that older adults may need to improve the ballistic capabilities of their arms to utilize them more effectively in the frontal plane to reduce center of mass dynamics and improve stability.

The current fall prevention programs could be missing another key element to create a more comprehensive protocol aimed at reducing falls in older adults. Some of the popular physical therapist led fall prevention programs like the Otago Exercise Program have been shown to be effective in reducing the fall rates in older adults (45). These programs focus on balancing and walking tasks, and heavy strengthening of the lower extremities at the ankle, knee and hips mainly focusing on flexion/extension strengthening (46,47). While these programs have shown success, it is possible that the decrements in arm movements exhibited by older adults are not being rehabilitated. Older adults’ reaction time to perturbations improve under training (48), and it is currently unknown if strength training of the upper extremities can improve kinematic responses. Improving functions of the arms that permit older adults to create ballistic upper extremity movements could increase the efficacy of fall prevention programs by making them more comprehensive.

The biomechanical function of the arms in older adults experiencing a slip should be further studied to better understand the mechanisms contributing to balance. A potential contributor to the higher fall rate in older adults experiencing the same arm motion as younger adults could be due to reported delays of shoulder movements in older adults exposed to a slip (27). A delay in mechanical response may allow the perturbation to continue its consequential effect on human body balance and make regaining balance more difficult. It is possible that musculoskeletal decrements in older adults lead to similar movements as younger adults, but at a slower velocity making their arm movements less effective on balance.

The results of this study should be interpreted through the lens of a number of limitations. The sample size of this study was quite small with only 11 older adults and 11 younger adults analyzed, and while this study would be strengthened with analyses of additional participants, it is still informative that the frontal plane arm motion was similar to previously reported studies utilizing younger adults. Lastly, our study allowed participants to wear their own shoes when stepping on a slippery surface whereas other studies standardize the shoes. This is important to consider as the shoe material, shoe treads, and the wear on the sole may affect slip severity. However, an important aspect to consider with this limitation is that people slip in the outside world wearing their own shoes in their own specific conditions.

## Summary

Older adults exhibit significantly lower arm acceleration and delayed arm abduction responses compared to younger adults in response to a slip incident. This resulted in older adults exhibiting significantly increased center of mass excursion which leads to a higher likelihood of losing balance. Improving the strength and ballistic capability of the arms in older adults can potentially lead to reduction in falls.

## Acknowledgements

We would like to thank Mark Grabiner, PhD for his contributions to this work.

